# Middle-way flexible docking: Pose prediction using mixed-resolution Monte Carlo in estrogen receptor *α*

**DOI:** 10.1101/424952

**Authors:** Justin Spiriti, Sundar Raman Subramanian, Rohith Palli, Maria Wu, Daniel M. Zuckerman

**Affiliations:** Department of Biomedical Engineering, Oregon Health and Science University, Portland, OR 97239; Address unknown; Medical Scientist Training Program and Biophysics, Structural & Computational Biology Program, University of Rochester, 601 Elmwood Avenue, Box 350, Rochester, NY 14642; 41521 Bloomin Spring Ct, Hebron KY 41048

## Abstract

There is a vast gulf between the two primary strategies for simulating protein-ligand interactions. Docking methods significantly limit or eliminate protein flexibility to gain great speed at the price of uncontrolled inaccuracy, whereas fully flexible atomistic molecular dynamics simulations are expensive and often suffer from limited sampling. We have developed a flexible docking approach geared especially for highly flexible or poorly resolved targets based on mixed-resolution Monte Carlo (MRMC), which is intended to offer a balance among speed, protein flexibility, and sampling power. The binding region of the protein is treated with a standard atomistic force field, while the remainder of the protein is modeled at the residue level with a Gō model that permits protein flexibility while saving computational cost. Implicit solvation is used. Here we assess three facets of the MRMC approach with implications for other docking studies: (i) the role of receptor flexibility in cross-docking pose prediction; (ii) the use of non-equilibrium candidate Monte Carlo (NCMC) and (iii) the use of pose-clustering in scoring. We examine 61 co-crystallized ligands of estrogen receptor *α*, an important cancer target known for its flexibility. We also compare the performance of the MRMC approach with Autodock smina, a docking program. [1] Adding protein flexibility, not surprisingly, leads to significantly lower total energies and stronger interactions between protein and ligand, but notably we document the important role of backbone flexibility in the improvement. The improved backbone flexibility also leads to improved performance relative to smina. Somewhat unexpectedly, our implementation of NCMC leads to only modestly improved sampling of ligand poses. Overall, the addition of protein flexibility improves the performance of docking, as measured by energy-ranked poses, but we do not find significant improvements based on cluster information or the use of NCMC.

## Introduction

Computational structure-based drug design can play an important role in drug development, as exemplified in the development of inhibitors of HIV protease, which have a major impact on treatment for AIDS patients. [2, 3] Because of their potential to reduce the cost and time associated with drug development, a multitude of methods have been developed to screen potential drug candidates virtually and prioritize possible structures for synthesis. [4–7] Some of the most popular docking methods represent the potential energy due to the receptor using grids and optimize the ligand conformation with respect to this potential. These include DOCK, [8, 9] Autodock Vina, [10] and the related smina [1], Schrodinger Glide, [11–13] CDOCKER, [14] and others. [15] Once a grid is constructed, however, the protein conformation represented by that grid is fixed, which is a serious approximation, since a number of structural studies have shown that “hidden” protein conformations and protein flexibility play important roles in protein-ligand binding; methods that consider protein flexibility and multiple structures improve docking performance compared to those that make use of only one structure. [16–23]

A number of approaches have been developed to incorporate flexibility into docking. [24] One approach is to try to incorporate protein flexibility into grid-based approaches by collecting multiple confomations of the receptor and docking to all of them, a practice commonly known as ensemble docking. [16,17,20,23,25,26] Another grid-based strategy involves leaving certain amino acid side chains out of the grid and optimizing their conformation alongside that of the ligand during the docking procedure. CDOCKER has been modified in this way for example, [27] and Autodock Vina and smina are also capable of this. [1, 10] However, the amount of flexibility that can be incorporated by this strategy is limited. In particular, allowing only a few side chains to be flexible means these few side chains must be carefully chosen and that important protein motions involving the backbone are not represented. Likewise, ensemble docking requires a careful choice of conformations to be used and only allows for a limited degree of backbone flexibility. Since these conformations come from simulations or structures of the protein without the corresponding ligand, ensemble docking cannot take into account the mutual induced fit that may occur when a ligand binds to a protein.

The RosettaLigand approach to ligand docking [28–30] makes use of the Rosetta knowledge-based force field and, in principle, allows for full receptor flexibility. Like other knowledge-based force fields, it relies on the assumption that the system under discussion is similar to known protein-ligand complexes. Furthermore, the RosettaLigand docking approach used a complex protocol involving several rounds of minimization, which loses information about the relative entropy of the energy minima that are found. Although in theory the protocol should be able to allow full receptor flexibility, in practice it was found that restraints on the *α* carbons were needed to improve the discrimination between native and nonnative poses. RosettaLigand is also relatively expensive computationally, requiring approximately 80 CPU hours per ligand. [28]

Another way in which full flexibility of the protein can be allowed, at even greater computational cost, is by using molecular dynamics or Monte Carlo simulations. Alchemical free energy methods have received much attention recently. [31–33] These methods require conducting multiple simulations at different values of a coupling parameter *λ* that serves to scale protein-ligand interactions. In principle, they are exact according to the laws of statistical mechanics and therefore should give perfectly accurate results given an accurate force field potential and a simulation of infinite length. In practice, however, both force field errors and inadequate sampling can result in significant errors in the calculated free energies. [34, 35] Alternatively, methods such as MM-PBSA [36, 37] can be used; these are somewhat less expensive because they only require simulation of the endpoints, but also make additional approximations. There is therefore a need for an *in silico* docking technique that incorporates full flexibility of the protein at modest computational cost.

Multiscale simulation techniques show promise for reducing computational time while maintaining full flexibility of the protein and physical accuracy. An early approach that bears some similarities our own involves dividing the system into three regions: an atomistic region, a coarse-grained region using a Gō-like potential, and an intermediate region. [38] Our group has also done some preliminary work on mixed resolution models, combining a residuel level Gō model with the OPLS-AA force field and conducting some preliminary tests on self-docking to the estrogen receptor. [39] The popular MARTINI force field for water has also been combined with an atomistic force field for proteins; [40] however, balancing the strength of electrostatic forces in the two force fields proved difficult. Others have tried to combine an atomistic region with an elastic network model. [41]. Feig and co-workers have also combined their PRIMO force field with the CHARMM36 atomistic force field, [42] and obtained similar results to fully atomistic or fully coarse-grained simulations, but they note issues with weakened hydrophobic packing interactions, and the amount of speedup they obtained is generally modest.

Here we extend our previous mixed-resolution Monte Carlo software [39] and use it to systematically study key aspects of docking in a challenging system. In the software, the majority of the protein is modeled using a residue-level coarse-grained model currently based on the Gō model [43–46] while the ligand and binding site are modeled using an atomistic force field. Interactions between the two regions are treated in a fully atomistic manner. Full flexibility of the ligand and receptor is maintained, allowing the modeling of mutual induced fit, including “breathing” motions away from the binding site. [47, 48] Importantly, the coordinates of all atoms are tracked throughout the simulation, which increases computational cost but enables maintaining proper backbone geometry via standard non-bonded terms even in the coarse region. Monte Carlo methods sample the Boltzmann distribution and hence implicitly include entropy effects. [49] The computational cost of the method can be adjusted by changing the size of the atomistic region; for the region used here, the mixed-resolution model reduces the computational cost by a factor of 2-4 compared to a fully atomistic treatment of the protein using the same implementation. In addition, coarse-grained models produce potential energy surfaces that are smoother than corresponding atomistic surfaces, because of averaging over omitted degrees of freedom. [50]

Monte Carlo simulation readily allows for the systematic testing of the role of flexibilty in docking, as has been noted. [28, 29] Flexible degrees of freedom and interaction type (coarse-grained vs. all-atom) are readily adjusted. Here, we specifically examine the effects of common choices in conventional docking: rigid side chains and rigid backbones, both of which prove detrimental. We are unaware of a prior study examining these cases together.

In addition to the mixed-resolution model, we examine the recently developed nonequilibrium candidate Monte Carlo (NCMC) method to enhance the sampling of ligand binding modes. [51] In this method, the potential energy is perturbed systematically in a short nonequilibrium simulation over the course of 10^2^-10^3^ MC trial moves, and the entire sequence of moves is then either accepted or rejected based on the nonequilibrium work done during the simulation. In principle, the perturbation of potential energy can be designed to allow large configurational changes that would have a very low acceptance probability in a standard MC simulation, while the short nonequilibrium simulations allow time for the system to relax after each such configurational change. Gill et al. have applied this method to the binding of toluene to the L99A mutant of T4 lysozyme and found that the NCMC method produced a much higher rate of transitions between distinct binding modes of toluene compared to standard MD simulations or simulations using MD combined with MC. [52]

We examine the effects of flexibility and the efficacy of NCMC in the context of docking known ligands to the ligand binding domain of estrogen receptor *α* (ER *α*). ER *α* is known to undergo a conformational change upon binding by estradiol and other ER agonists in which helix 12 closes over the binding site as shown in Fig 1. [53, 54] In contrast, in the bound structures with tamoxifen and other ER *α* antagonists, helix 12 does not make this conformational change. [55] The development of drugs to modulate ER activity is of considerable interest because aberrant ER signaling has long been known to be a key player in promoting proliferation of several types of cancer, and multiple ER modulating drugs are currently in clinical use. [56–58] However ER has also been recently observed to evolve drug-resistance mutations during metastatic progression in breast cancer, which limits the ability of current therapeutic agents to affect the progression of secondary tumors. [59–61] We have previously validated a similar MRMC approach for ER in a very limited way using a simple self-docking test. [39] Here we attack the cross-docking problem, in which the co-crystal structure for the tested compound is not used.

**Fig 1.**
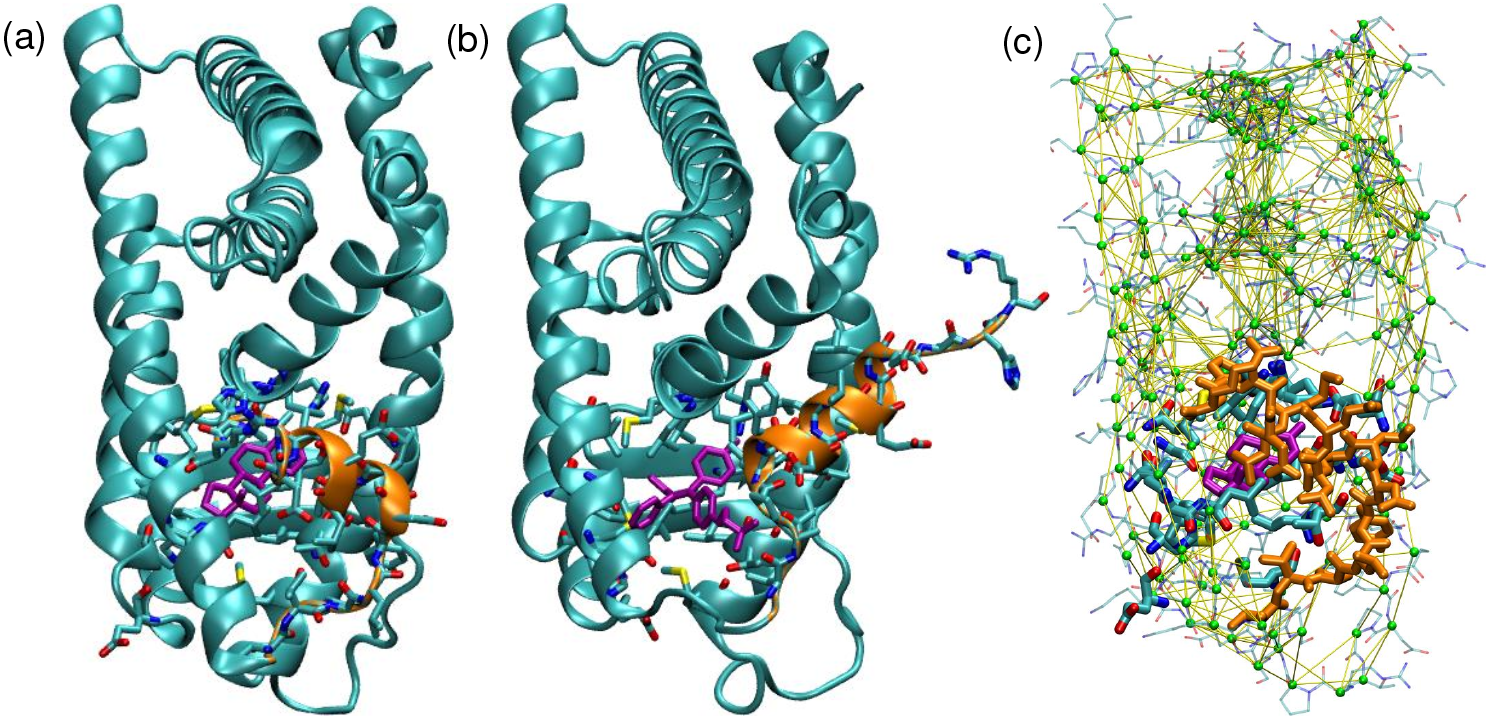
Structure of ER *α*. (a) Active and (b) inactive conformation of ER *α*. (c) Illustration of the mixed-resolution model used in this paper. The ligand (green) and the binding site (heavy structure) constitute the atomistic region and are treated using an all-atom force field. The remainder of the protein is treated as the coarse-grained region is represented by particles located at each *α* carbon (purple spheres) with native attractions between them (yellow lines). Rigid structures of each amino acid (thin structure) and moved along with the coarse-grained particles and used to calculate interactions between the coarse-grained and atomistic regions. Helix 12 is indicated in orange in all three panels.

The first section of the paper describes the mixed resolution potential and the Monte Carlo and NCMC protocols that were used with it for docking. For this study, ligands with known bound crystal structures were used, so we can compare the docked conformations to experimental crystal structures. We study the docking protocol with and without NCMC and with different levels of protein flexibility, and also tested different methods of ranking the poses, including trying to cluster the poses by structural similarity. We find that having full protein flexibility (including backbone flexibility) results in stronger interactions between protein and ligand, and top-ranked poses that are closer to the corresponding crystal structures. Study of the acceptance rates of large ligand moves and of variations in ligand RMSD during each simulation provide some evidence that NCMC is improving sampling. However, the use of NCMC does not improve the docking results beyond what is obtained with simulations with a fully flexible protein without NCMC. Also, the use of clustering as a part of pose ranking does not appear to improve the overall docking results. We also compare the performance of the MRMC method to Autodock smina, [1] chosen to represent docking software, and find that the increased backbone flexibility offered by MRMC improves performance compared to smina. Finally, we discuss possible improvements to the mixed resolution potential and to sampling, and the ability of our protocol to sample multiple docked poses for each ligand.

## Materials and methods

### Mixed-Resolution Potential

In this work, we have replaced the discontinous Gō model functional form used in previous work [45, 46] with a continous Lennard-Jones functional form. We have also replaced the OPLS-AA force field used in our previous work with the AMBER 99SB force field [62] so that ligands can be automatically parameterized using the compatible GAFF forcefield and antechamber tool. [63, 64] To save computer time, we have replaced the Generalized Born solvent model with a simpler solvent exposure-dependent distance-dependent dielectric model. [65] We no longer use precalculated libraries of amino acid conformations in our Monte Carlo moves—a technique that worked well for peptides but proved to be less advantageous for Monte Carlo simulations of dense protein systems because of poor acceptance rates.

Simulations were done with a mixed-resolution potential, in which the majority of the protein is treated with a Gō model [43–46, 66] and the atomistic region around the binding site is treated using the AMBER 99SB force field. [62] The overall potential has the following form:

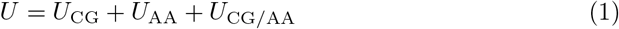

The coarse-grained portion of the potential *U*_CG_ is in turn given by

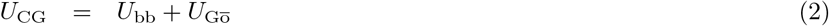

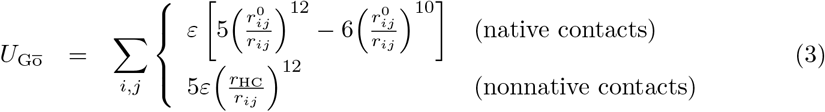

where the sum is taken over all pairs of residues in which both residues belong to the coarse-grained region. The coarse-grained potential also includes a backbone component, which uses the bond, angle, and dihedral terms from the AMBER 99SB force field for the backbone atoms in the coarse-grained region (1-4 van der Waals terms are not included). The 12-10 form used here replaces a square well potential used in previous work [45, 46] and has shown good performance in other settings. [66–68] The Gō model well depth *ε* and hard core radius *r_HC_* are listed in table 1. 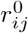 is the native distance between atoms *i* and *j*, determined from the original structure.

**Table 1.** Parameters of the simulation.

| Symbol | Description | Value |
| --- | --- | --- |
| $\varepsilon$ | Gō model well depth | 3.0 kcal/mol |
| $r_{\text{HC}}$ | Hard core radius | 1.7 Å |
|  | Cutoff for defining native interactions in Gō model | 8 Å |
| $\varepsilon_0$ | Low-dielectric constant | 2 |
| $\varepsilon_1$ | High dielectric constant | 8 |
| $c$ | Constant for determining hydration overlaps | 0.625 |
|  | Cutoff for atomistic nonbonded interactions | 10 Å |
| $T$ | MC simulation temperature | 300 K |
|  | Method of selecting atomistic region | residues 533-548 plus all residues with at least one non-hydrogen atom within 3 Å of the ligand in any of 66 reference structures |
|  | Number of residues in atomistic region | 39 |

The all-atom potential *U*_AA_ is the potential energy of the all-atom region, according to the AMBER 99SB force field. [62] The *U*_CG/AA_ term represents the interaction between the two regions, which is also computed atomistically using the AMBER 99SB force field. The all-atom region was defined to contain the ligand and all residues with at least one non-hydrogen atom within 3Å of the ligand in any of the 66 crystal structures of ligand-ER complexes that were used as references; in addition, residues 533-548, which comprise helix 12 and the neighboring loop, were also included in the all-atom region. Ligands were parameterized with the antechamber tool [64] using the GAFF force field. [63]

In order to incorporate solvation effects in a computationally efficient manner, the electrostatic term of the AMBER 99SB was modified to use a solvent-exposure dependent distance dependent dielectric originally developed by Garden and Zhorov, which was previously found to work well in docking simulations with an AMBER force field. [65] In this model, the electrostatic interaction between atoms *i* and *j* is given by *K_coul_q_i_q_j_/ε_ij_r_ij_* where

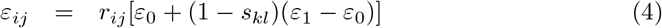

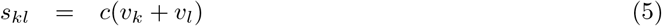

where *ε*_0_ and *ε*_1_ are low and high dielectric constants, respectively, and *s_kl_* is the overlap of hydration shell volumes for the groups *k* and *l* that include atoms *i* and *j*. (For all pairs of groups *k* and l, 0 ≤ *s_kl_* < 1.) The hydration volumes *v_k_* and *v_l_* for these groups are calculated using a formula originally used for the EEF1 implicit solvent model. [69] The values for *ε*_0_, *ε*_1_, and *c* shown in table 1 are those found to optimize docking results for a training set in ref. 65.

### Standard Monte Carlo

Sampling was performed using MC simulation employing a standard variety of local and global moves as described in table 2. To investigate the impact of protein flexiblity, in addition to simulations in which the protein was fully flexible, simulations were also conducted in which the protein was kept rigid or only the sidechains were allowed to move. This was done by leaving out the corresponding MC moves and increasing the fraction of others in the move mix, also as shown in table 2. In all simulations, both the translational and rotational degrees of freedom of the ligand with respect to the protein and its internal degrees of freedom were sampled.

**Table 2.**
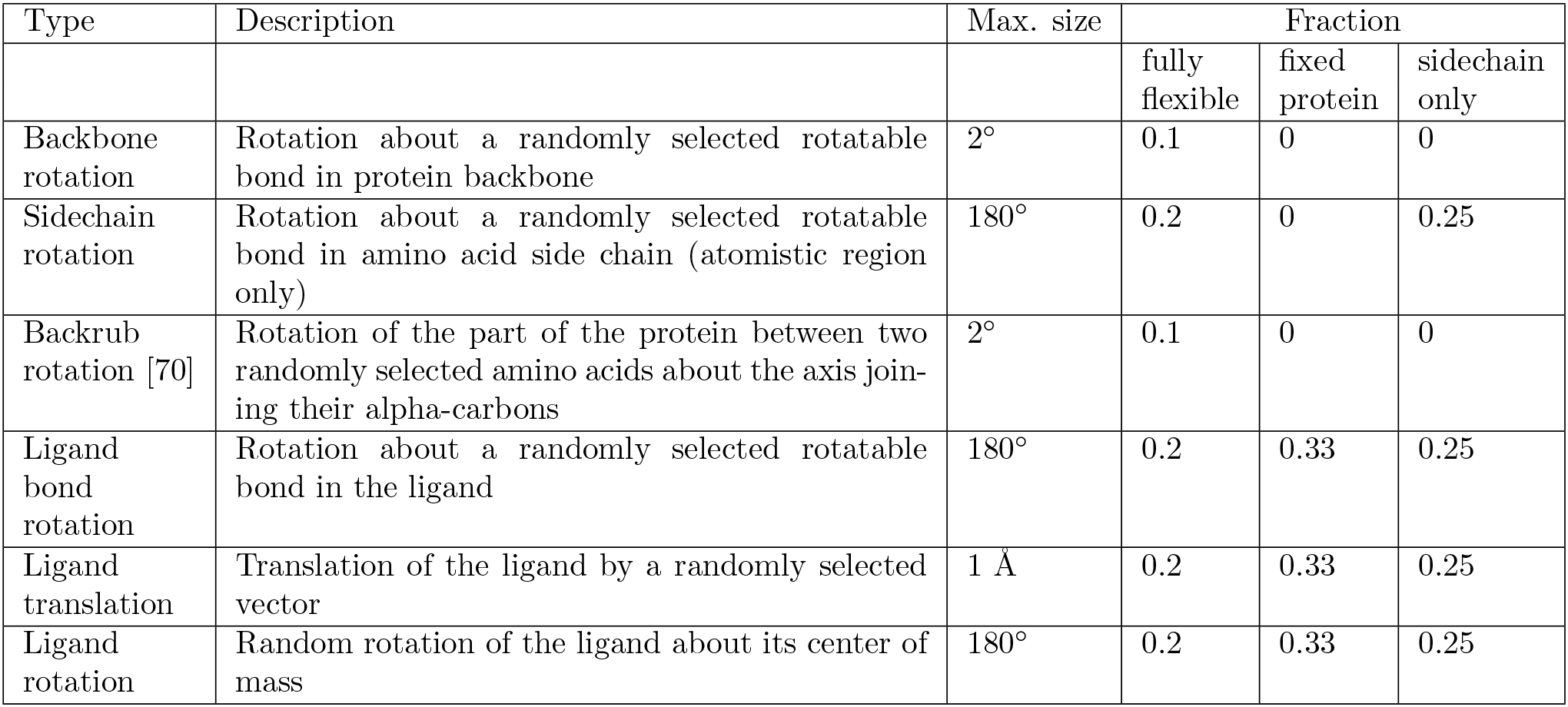
Monte Carlo moves used in the simulation.

| Type | Description | Max. size | Fraction |  |  |
| --- | --- | --- | --- | --- | --- |
|  |  |  | fully flexible | fixed protein | sidechain only |
| Backbone rotation | Rotation about a randomly selected rotatable bond in protein backbone | 2° | 0.1 | 0 | 0 |
| Sidechain rotation | Rotation about a randomly selected rotatable bond in amino acid side chain (atomistic region only) | 180° | 0.2 | 0 | 0.25 |
| Backrub rotation [70] | Rotation of the part of the protein between two randomly selected amino acids about the axis joining their alpha-carbons | 2° | 0.1 | 0 | 0 |
| Ligand bond rotation | Rotation about a randomly selected rotatable bond in the ligand | 180° | 0.2 | 0.33 | 0.25 |
| Ligand translation | Translation of the ligand by a randomly selected vector | 1 Å | 0.2 | 0.33 | 0.25 |
| Ligand rotation | Random rotation of the ligand about its center of mass | 180° | 0.2 | 0.33 | 0.25 |

### Nonequilibrium candidate Monte Carlo

In order to enhance the sampling of ligand poses, docking runs were also undertaken using a modified version of the nonequilibrium candidate Monte Carlo (NCMC) algorithm. [51, 52] In this method, the system is subjected to NCMC “moves” which are in fact short nonequilibrium simulations (using standard MC) during which the potential energy function is occasionally perturbed such that sampling for particular degrees of freedom may be enhanced.

In the method used here, the potential energy is modified by scaling the van der Waals and electrostatic components of the protein-ligand interaction energy by the factors *λ*_VDW_ and *λ*_elec_:

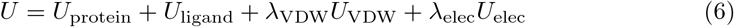

Each NCMC move consisted of 800 individual Monte Carlo moves, which were divided into four 100-move phases in which the coupling parameters *λ*_VDW_ and *λ*_elec_ were changed according to the schedule shown in Fig. 2. First, the charges on the ligand were removed by systematically driving *λ*_elec_ towards 0. Second, *λ*_VDW_ was also driven towards 0, such that the ligand was completely uncoupled from the protein and free to rotate, translate, or change conformation without interference. Then, in the second two phases, first *λ*_VDW_ and then *λ*_elec_ weree gradually transitioned back to 1, so that the system relaxed and steric clashes might be resolved. In all four phases, *λ*_VDW_ and *λ*_elec_ were changed by 0.05 every 5 trial moves. During all four phases, the individual trial moves were preliminarily accepted or rejected with the standard Metropolis criterion,

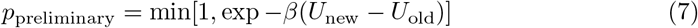

where *U* is the scaled potential given above. This leads to canonical sampling suitable for the given set of *λ* values.

**Fig 2.**
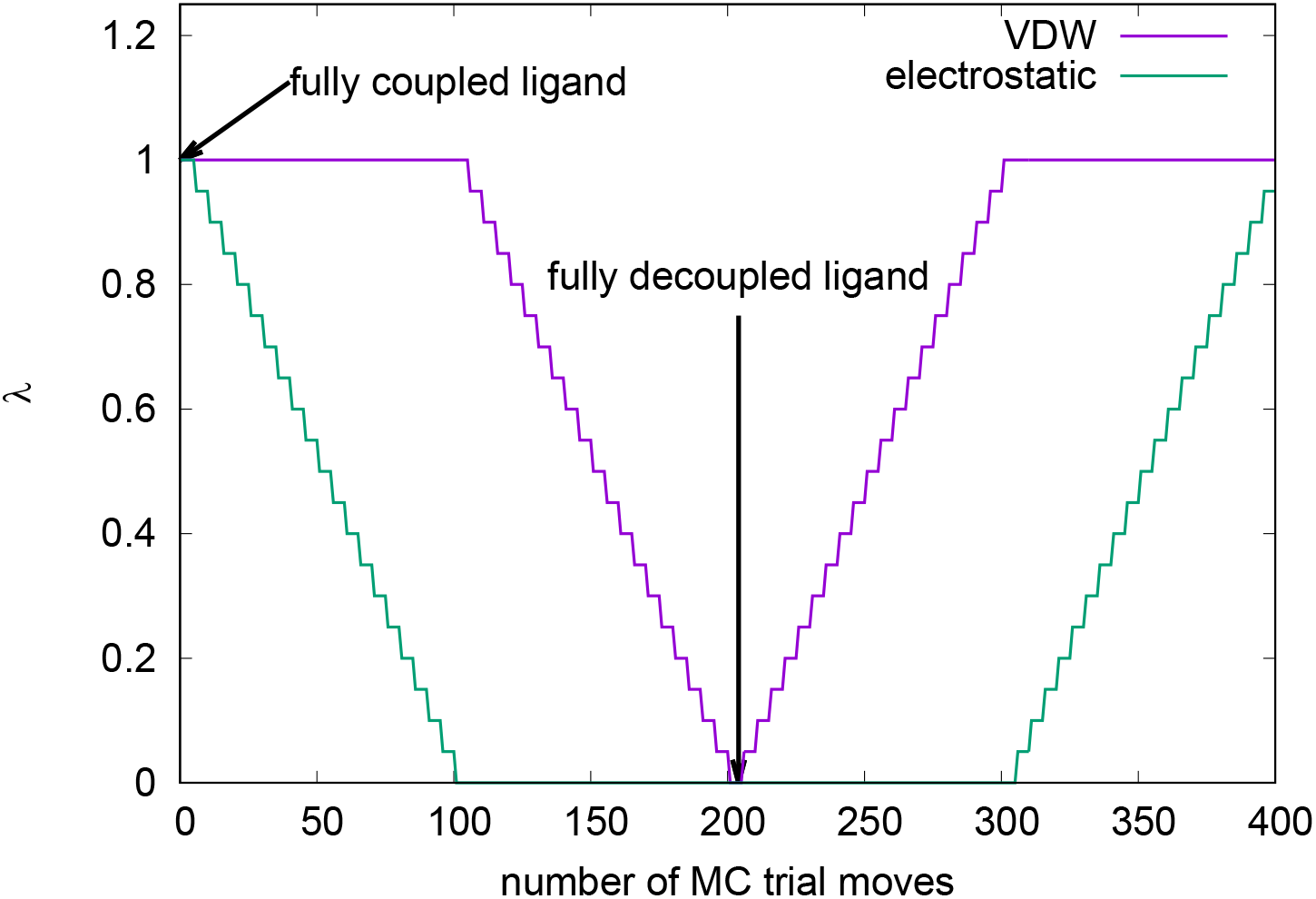
Scaling of *λ*_VDW_ and *λ*_elec_ over each NCMC trial-move cycle. The cycle comprises four phases, each containing 100 individual MC trial moves. First, the charges on the ligand are removed by gradually reducing *λ*_elec_ to 0. Second, the ligand is fully uncoupled from the protein by reducing *λ*_VDW_ to 0. Third, the van der Waals interactions of the ligand with the protein are restored by increasing *λ*_VDW_ back to 1. Finally, the charges on the ligand are restored by increasing *λ*_elec_ back to 1.

At the end of each NCMC move, the nonequilibrium work *w* performed on the system (which accounts for the changes in *λ* values) was calculated using

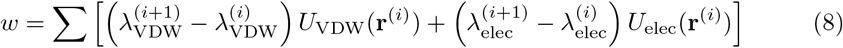

In this equation, the index *i* enumerates values of *λ*_VDW_ or *λ*_elec_ that are used during the move (horizontal segments of the graph in Fig. 2) and there is a term in the sum for each change in the *λ* parameters. The full sequence of MC moves that had taken place during the MC trial was then finally accepted or rejected with probability given by

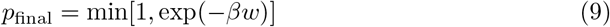

If the NCMC move was rejected, the conformation of the system prior to the sequence of MC steps making up the given NCMC move was restored. Because of this, and the low acceptance rate of NCMC moves (about 2%), in pure NCMC simulations the relaxation of the system toward low energy conformations was extremely slow. Consequently, in order to promote more rapid relaxation, the NCMC moves were alternated with 400 moves of regular MC.

### Docking protocol

A total of 61 ligands were used in this work, 37 agonists and 24 antagonists. Each ligand had a corresponding reference crystal structure, drawn from the PDB, showing the experimentally determined bound structure. Lists of the ligands and their corresponding reference structures are found in tables S1 and S2. Figure 3 shows example ligand structures, where the larger size and flexibility of antagonists is notable.

**Fig 3.**
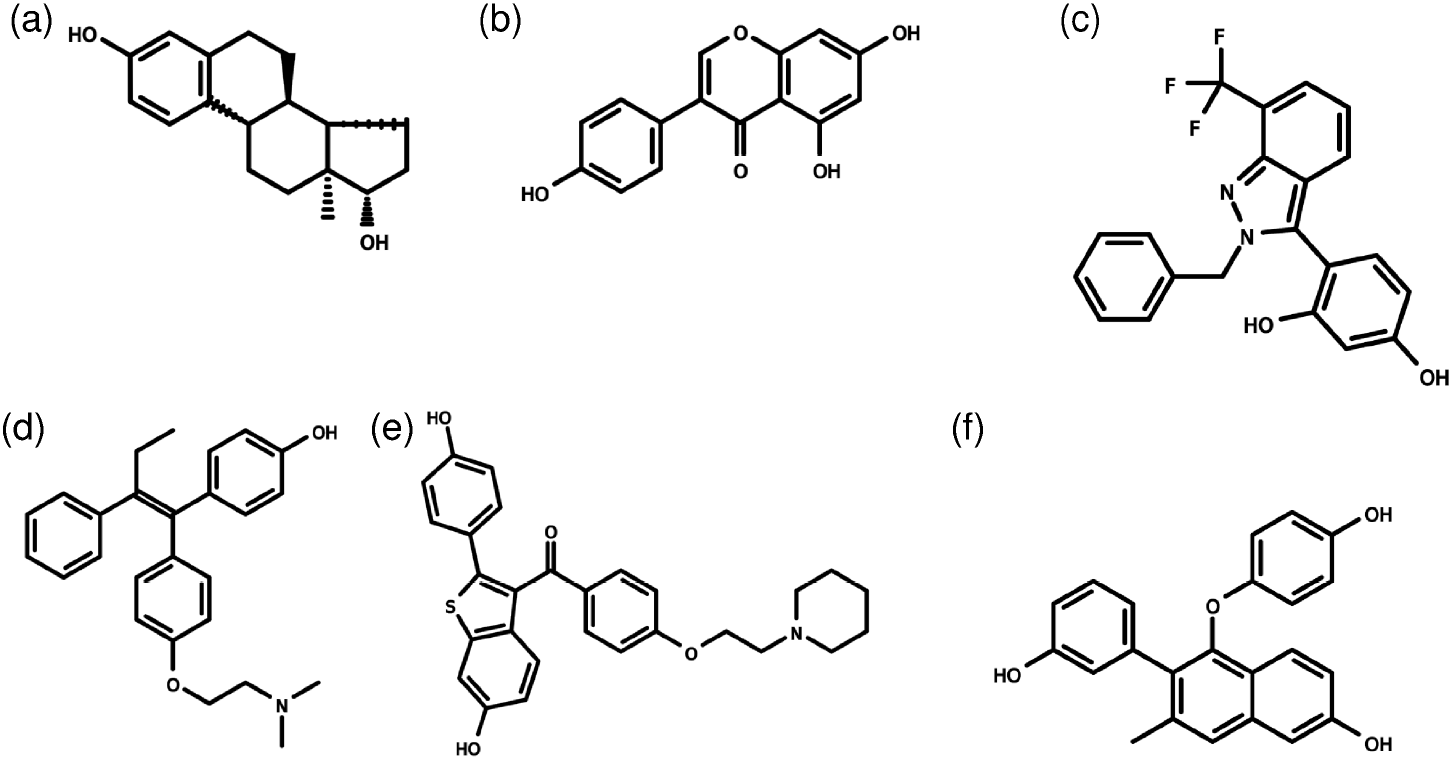
Examples of ligands studied in this work. (a) estradiol; (b) genistein; (c) the drug 1GJ; (d) 4-hydroxytamoxifen; (e) raloxifene; (f) the drug 369. (a)-(c) are agonists, whereas (d)-(f) are antagonists.

Depending on whether the compound in question was an agonist or antagonist, it was cross-docked against either the active or inactive conformation of ER. (Although in principle the MRMC method is capable of simulating the transition between the active and inactive conformations of ER, such a simulation would likely be longer than the docking runs used here, so docking of agonists against the inactive conformation or of antagonists against the active conformation was not attempted.) The active conformation was taken from the crystal structure of ER in complex with estradiol (PDB code 1QKU) [54] whereas the inactive conformation was taken from the crystal structure in complex with 4-hydroxytamoxifen (PDB code 3ERT). [55] Hydrogen atoms were added using the tleap tool in AMBER, and ionization states for the twelve histidine residues were chosen based on a combination of pKa calculations made using H++ [71, 72] and visual inspection of the crystal structures. Histidine residues 356, 373, 398, 476, 488, 501, 513, and 516 were chosen to be neutral, while histidine residues 377, 474, and 547 were chosen to be ionized. In the active conformation, His 524 makes a hydrogen bond to the hydroxyl group on C17 of estradiol, and it was calcuated to have a pKa of 5.67 using H++, implying a neutral state. In contrast, in the inactive conformation, His 524 makes a salt bridge with Glu 419 and was found to have a pKa of 7.97, implying an ionized state. Given the ambiguous nature of His 524’s ionization state and its potential importance for the accuracy of docking results, it was decided to conduct half of the docking runs with an ionized His 524 and half with a neutral His 524.

Structure data files for each ligand were downloaded in SDF format from the Protein Data Bank and used to generate initial coordinates for each ligand. As a part of this process, ionization states for each ligand at pH 7 were determined using OpenBabel [73] and hydrogen atoms were added accordingly.

Fig. 4 gives an overview of the docking and analysis procedure. Each docking run consisted of a search for initial low-energy poses followed by refinement using Monte Carlo simulation. For the initial search, 1000 random poses were generated by placing the ligand in a random position and orientation within 4 A of the center of mass of estradiol (for agonists) or 4-hydroxytamoxifen (for antagonists) in the crystal structures and rotating every rotatable bond in the ligand through a random angle. The energy of each of these random poses was computed and a Monte Carlo simulation lasting either 40000 trial moves (for regular MC) or 80000 trial moves (for mixed NCMC/MC) was started from the lowest energy pose. A total of 60 docking runs were performed with each His 524 ionization state as described above, for a total of 120 docking runs per drug overall. Each docking run took 2-4 hours on a single CPU, for a total of approximately 240 to 480 CPU-hours per drug.

**Fig 4.**
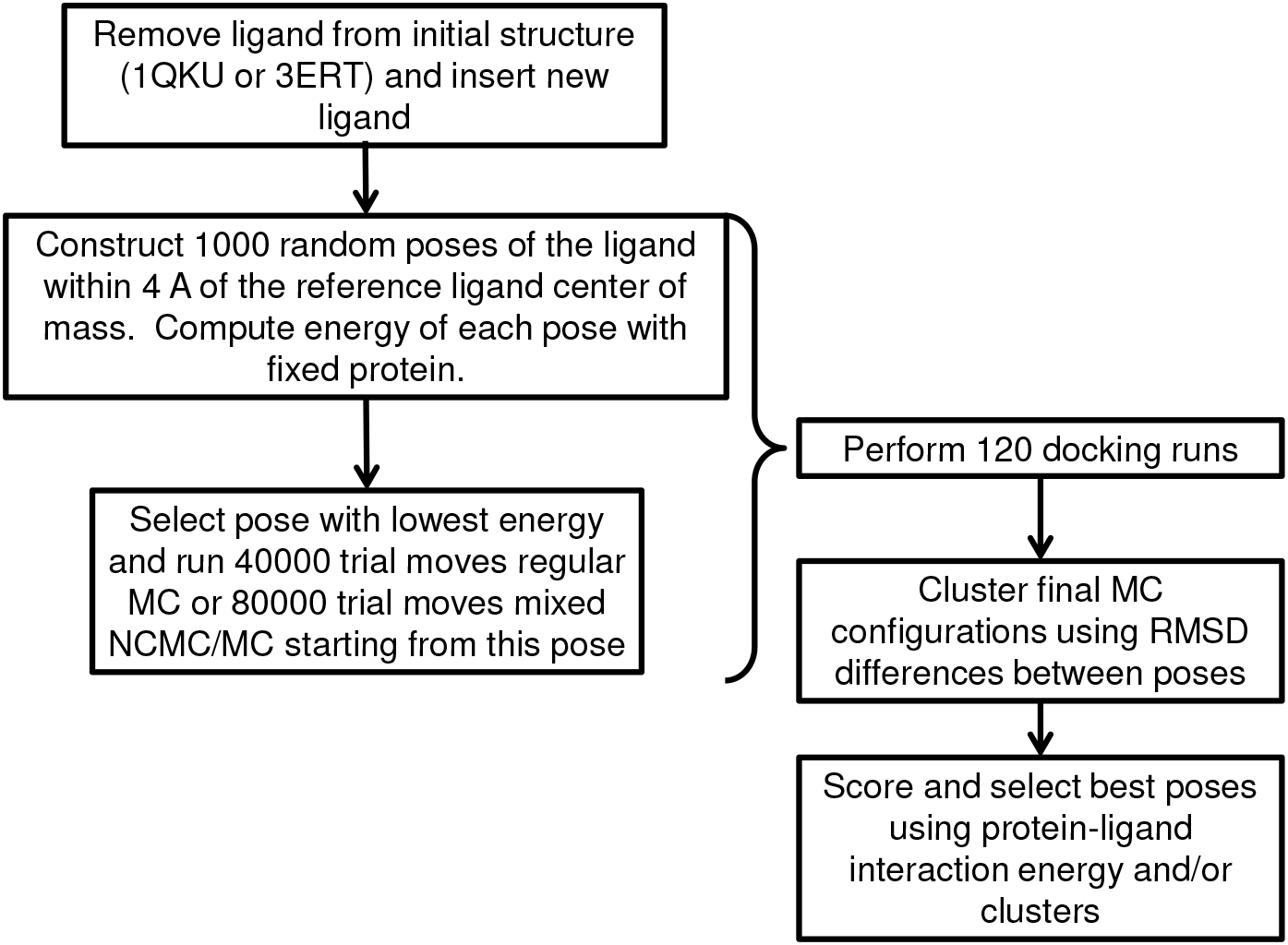
Flowchart showing overview of docking, clustering and scoring procedures used in this paper.

### Structural Analysis and Clustering

An important measure of the performance of a docking method is the heavy-atom RMSD of the ligand in the docked pose to a known crystal pose. These measurements could be made for all the ligands tested in this work, since corresponding crystal structures were available. Ligand RMSDs were measured with the protein backbone aligned to the reference structure, and taking into account all possible mappings of chemically equivalent groups on the ligand. The measurements were made using VMD. [74]

Clustering of structures was accomplished by first aligning the structures for each ligand according to the protein backbone and then constructing a pairwise distance matrix among them using ligand heavy atom RMSD as the metric. Complete linkage hierarchical clustering [75] was then applied to this distance matrix to construct a “phylogenetic tree” using ligand heavy atom RMSD taking into account chemically equivalent groups. The clusters were then defined from this tree, using a cutoff defined as the 10th percentile of all distances between structures for each drug. This ensured that the size of the clusters in configuration space would be appropriately scaled to the overall distribution of the poses for each drug.

### Comparison with other docking programs

In order to compare the performance of MRMC to other docking programs, we also performed docking with smina [1]. We carried out docking runs with smina using the default energy function and weights, either with a rigid protein or allowing flexible side chains for all of the amino acids that were in the atomistic region in the MRMC docking runs. One run of smina was performed with each level of flexibility for each drug and His524 titration state. Energy cutoffs were set to high values in order to recover as many docked poses as possible from each run, although the number of poses was limited to 60 so as not to exceed the number obtained from the MRMC docking runs. The docking runs were carried out in parallel using 16 CPUs at a time. For each pose, the heavy-atom RMSD was calculated relative to the crystal pose in the same manner as for the MRMC runs, and summary information was compiled in the same manner as for MRMC.

## Results

### Impact of Protein Flexibility

We first examined the impact of flexibility on pose generation. Fig.shows scatterplots of ligand RMSD versus interaction energy for two representative agonists (estradiol and genistein) and two representative antagonists (4-hydroxytamoxifen and raloxifene). Since the starting structures used were those of ER *α* bound to estradiol or 4-hydroxytamoxifen, the simulations with these ligands represent redocking, whereas those with all other ligands represent cross-docking. In fig., simulations in which the protein is fully flexible are compared with simulations in which the protein is rigid, or in which only the sidechains are allowed to move. Full protein flexibility results in interaction energies that are lower than those obtained with flexible side chains, which are in turn lower than those obtained with a completely fixed protein. In some cases - e.g. Panel (d) - the lack of flexibility prevents discovery of low-RMSD poses. This demonstrates that the use of protein flexibility, and particularly backbone flexibility, results in final configurations with stronger interactions between protein and ligand; this point is quantified further below.

The distribution of RMSDs suggests that the final configurations can be divided into clusters in some cases. This is particularly true for the agonists estradiol and genistein, which are relatively flat, rigid drugs and can fit into the binding site in multiple orientations. In these cases, the clusters correspond to these distinct orientations of the drugs. The ER antagonists 4-hydroxytamoxifen and raloxifene are more flexible; consequently, the clusters due to binding in multiple orientations are less clear.

Fig. 5 also shows a comparison between regular Monte Carlo simulations and those conducted with the mixed NCMC/MC protocol. The mixed NCMC/MC simulations had slightly higher interaction energies than the regular MC simulations.

**Fig 5.**
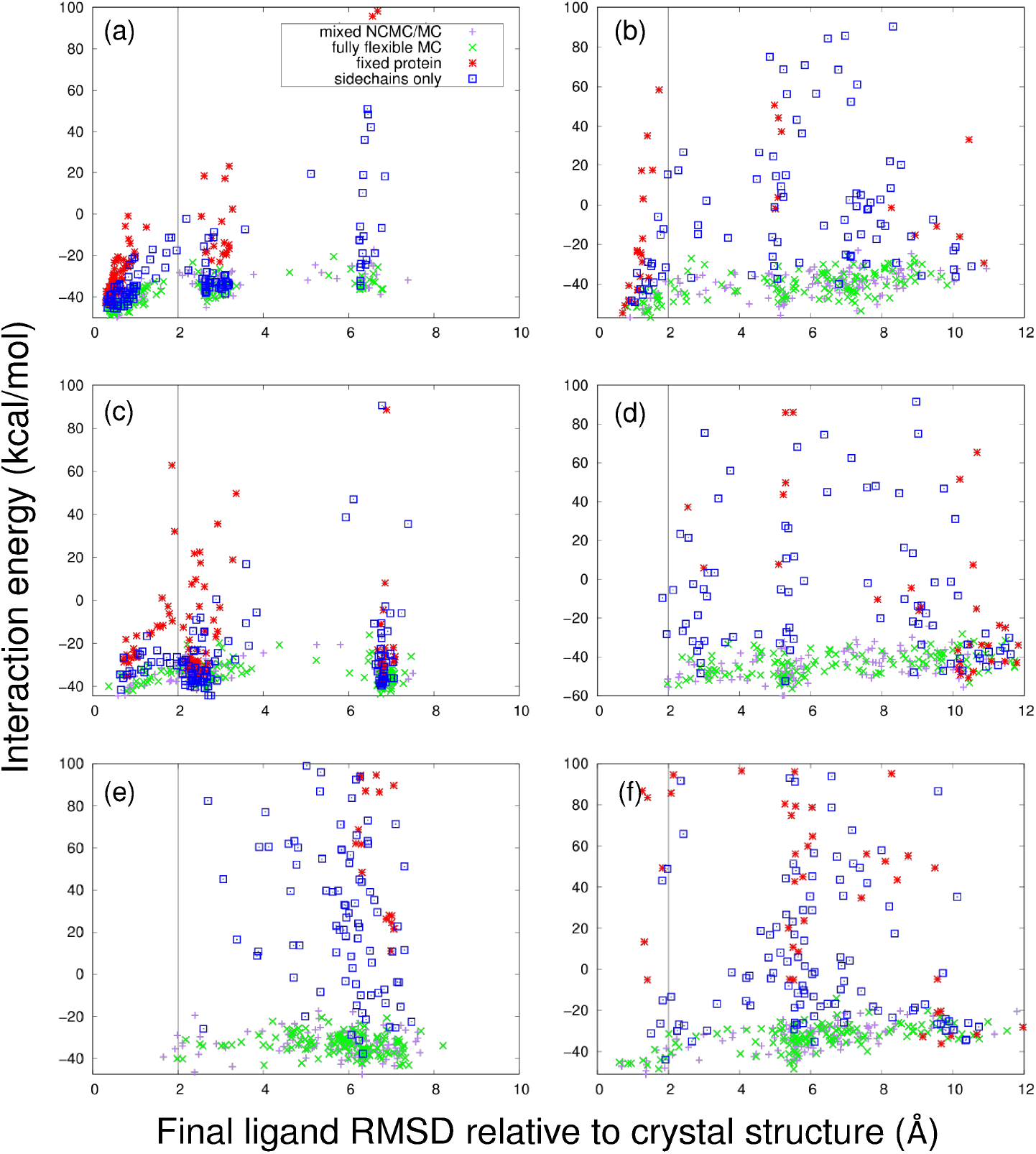
Example ensemble redocking and cross-docking runs for simulations incorporating different levels of flexibility. Plot of final ligand RMSD relative to crystal structure vs. interaction energy for (a) estradiol (redocking); (b) 4-hydroxytamoxifen (redocking); (c) genistein (cross-docking); (d) raloxifene (cross-docking); (e) the drug 1GJ; (f) the drug 369. (For structures of the drugs, see fig. 3.) (a), (c), and (e) are agonists, whereas (b), (d), and (f) are antagonists. In each plot, results for a fully flexible protein are in green (for MC only) or purple (for the mixed NCMC/MC simulations), whereas results for docking simulations in which the entire protein or just its backbone are fixed (MC only simulations) are in red or blue respectively. In order to make the results for the fully flexible protein visible, the vertical axis for each plot is cut off at 100 kcal/mol; as a result, some of the docking runs for the fixed protein are not shown due to their high energies.

The effect of protein flexibility is also demonstrated by a comparison of the “best” poses (selected on the basis of protein-ligand interaction energy) generated by our protocol, shown in Fig. 6. Figs. 6a and 6b show the distribution of “best” pose RMSDs across all of the drugs, expressed as a cumulative density function. Figs. 6c and 6d show a different approach to evaluating our protocol. For each drug, the best *N* poses are selected on the basis of protein-ligand interaction energy, the minimum RMSD pose is selected as a function of these, and the RMSD is averaged over all drugs. This average RMSD is then plotted as a function of *N*. It is clear that a much greater proportion of these best poses are close to the corresponding crystal structures when the protein is allowed to be fully flexible than when the protein is rigid or only sidechains move, and that the average best RMSD achieved is lower as well when more flexibility is allowed. This is especially true for the antagonists, which are generally more flexible than the agonists and whose docking is consequently more challenging. This difference is probably because the inactive structure of ER *α* is more open and consequently the protein as a whole, particularly helix 12, is more flexible. Consequently, the additional flexibility provided by the MRMC method may be more important for the inactive state.

In principle, allowing protein flexibility should enable us also to predict the changes in protein structure that occur upon docking to different ligands. We constructed Ramachandran and Janin plots [76] for the amino acid residues in the atomistic region for the final conformations from our docking runs. Fig. 7 shows these plots for two representative compounds, genistein and raloxifene. Although the overall conformation of ER around the active site is very similar for all bound agonists (and likewise for antagonists) there are slight differences in the backbone conformation for different ligands. The Janin plots (Figs. 7b and 7d) show relatively thorough sampling of free energy basins in the *χ_1_ -χ_2_* plane. The Ramachandran plots (Figs. (Figs. 7a and 7c), on the other hand, show that the backbone sampling is very limited (at least in the binding region) and that final conformations are remaining close to the initial structures rather than adapting to the different ligands. Nevertheless, the overall backbone flexibility which includes the CG region evidently has a significant effect on the generated poses, as shown in fig. 6.

**Fig 6.**
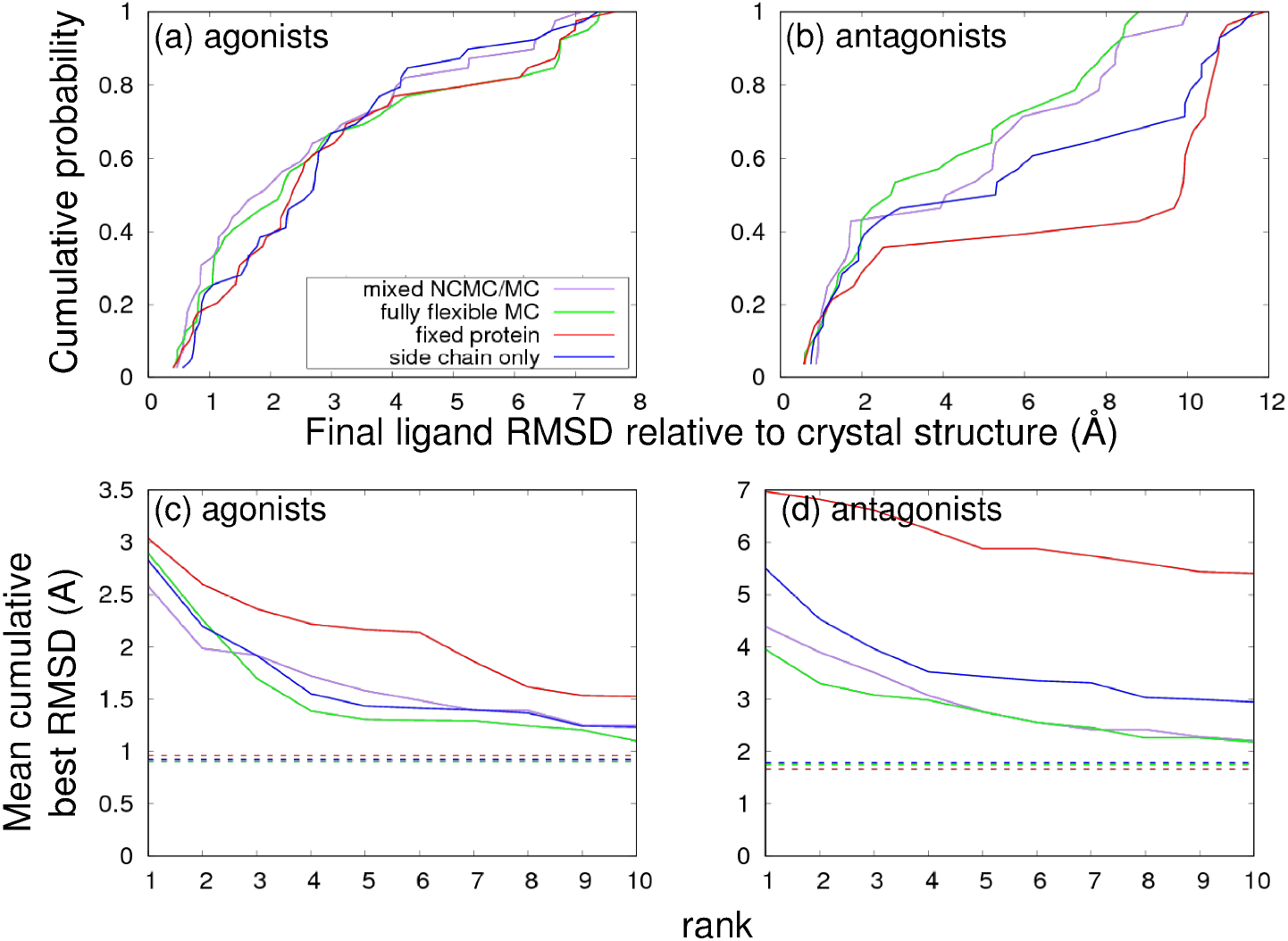
Performance of docking protocols. (a) and (b) show the cumulative probability distribution of RMSD values for the docked poses with the lowest final interaction energies for each drug, for (a) agonists or (b) antagonists. Larger values at low RMSD indicate better performance. In (c) and (d), for each drug the docking runs with the *N* lowest interaction energies are chosen and the best RMSD from among these is averaged across (c) agonists or (d) antagonists, so that lower RMSD indicates better performance. This average RMSD is plotted against *N*; dashed horizontal lines indicate the average best RMSD overall, without regard for interaction energy.

**Fig 7.**
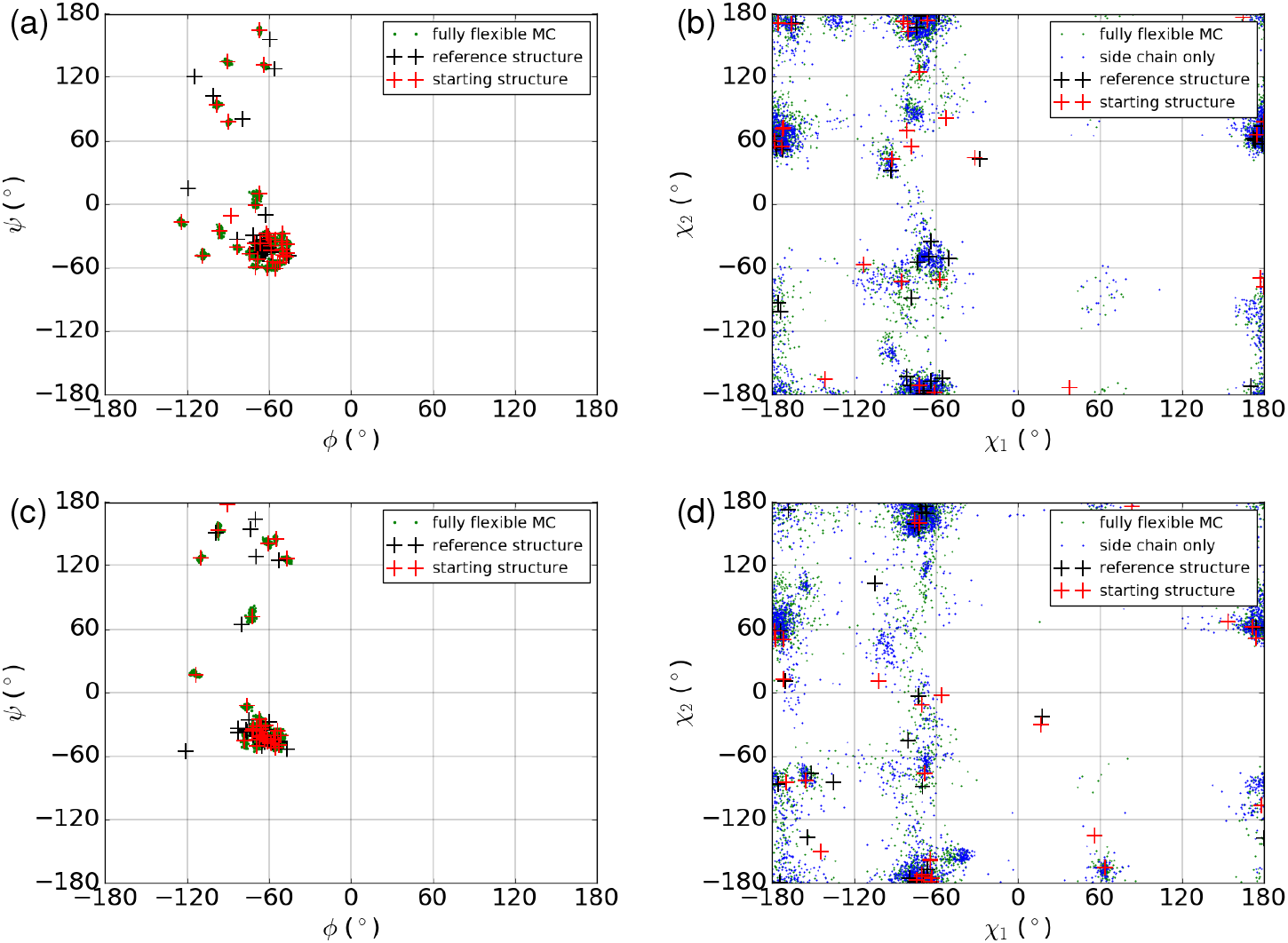
Examination of backbone and sidechain flexibility for two compounds. Ramachandran and Janin (*χ*_1_ vs. *χ*_2_) plots are shown for for final conformations from docking runs for (a)-(b) genistein and (c)-(d) raloxifene. Plots include only amino acids in the atomistic region. The reference structure is the corresponding crystal structure for each cross-docked compound.

### Assessment of NCMC

Fig. 6 also compares the performance of our docking protocol with and without NCMC. The use of NCMC shows at best a modest improvement in the overall docking performance, indicating that the NCMC is not enhancing sampling as much as was expected. In order to investigate this further, we measured both the distribution of overall ligand move sizes generated as well as the average acceptance probability as a function of move size. In the case of the NCMC simulations the size of the overall NCMC move depended on the individual MC moves that were performed within that move. To assess these factors, for each NCMC move, the overall rotation and translation of the ligand in the frame of reference of the protein was determined by first performing an RMSD alignment of the protein backbone of the final configuration relative to the initial configuration, then measuring the overall translation and rotation of the ligands needed to minimize the RMSD of the ligand heavy atoms in the two configurations. The cumulative density function of the overall magnitude of the displacement or the overall angle of rotation, and the average final acceptance probability (given by eq. 9) as a function of the overall displacement or rotation angle, were both computed and plotted as shown in Fig. S1. The corresponding distribution of move sizes and average acceptance probability were also plotted for comparison.

The plots show that, compared to regular MC, the distribution of moves generated by NCMC favors smaller moves in both translation and rotation. In addition, while acceptance rates for small moves are comparable for NCMC and MC, the acceptance rates for larger moves are many orders of magnitude smaller for NCMC. This is particularly (and unexpectedly) true for ligand rotations, where acceptance rates for all but the smallest rotations are much smaller for NCMC than for MC. The combined effect of these two trends is that in NCMC, a greater number of small translations and rotations are generated and accepted, compared to MC. The reason for these results seems to be that large ligand translations and rotations typically place the ligand in positions that clash sterically with the protein. Relaxing away such clashes evidently requires large nonequilibrium work and consequently leads to rejections. We note that these results are specific to our implementation, as described below in the Discussion.

Fig. S2 shows time series of the ligand RMSD for a number of individual docking simulations in both regular MC and NCMC. In many of the simulations, large jumps in RMSD can be seen; these represent large ligand moves that have been accepted. While the docking runs with the mixed NCMC/MC protocol are twice as long as those with regular MC, more than twice as many RMSD jumps can be seen in trajectories using the mixed NCMC/MC protocol. To reconcile this result with the acceptance rate data shown in Fig. S1, note that some of the NCMC transitions occur via a large number of small moves. It should also be noted that NCMC moves also include moving the protein, whereas individual MC moves that involve ligand translation and rotation do not, so Fig. S2 is not a perfect comparison.

### Comparison with Autodock smina

To assess the value of the MRMC approach compared with conventional docking software, we studied the same set of ligands and receptor structures using Autodock smina, both with the protein fixed and with all the side chains in the atomistic region allowed to move. [1] As shown in fig. 8, when used with full flexibility, MRMC generally outperformed Autodock smina, and particularly so for agonists, due primarily to the additional flexibility MRMC offers. (The cumulative distribution function of pose RMSD for antagonists shows somewhat better performance for smina with a fixed protein compared to MRMC, but this is not confirmed by the study of average RMSDs.) It is also of interest to compare the performance of MRMC to smina when both are used with the same amount of protein flexibility. It appears that MRMC performs better than smina for agonists, whether the protein is fixed or side chains within the MRMC all-atom region are allowed to move. The situation is more ambiguous for antagonists. There are several possible reasons for these differences, including differences in the treatment of solvation and salt bridge interactions that are crucial to determining the relative orientation of agonists within the ER active site, as well as differences in sampling.

**Fig 8.**
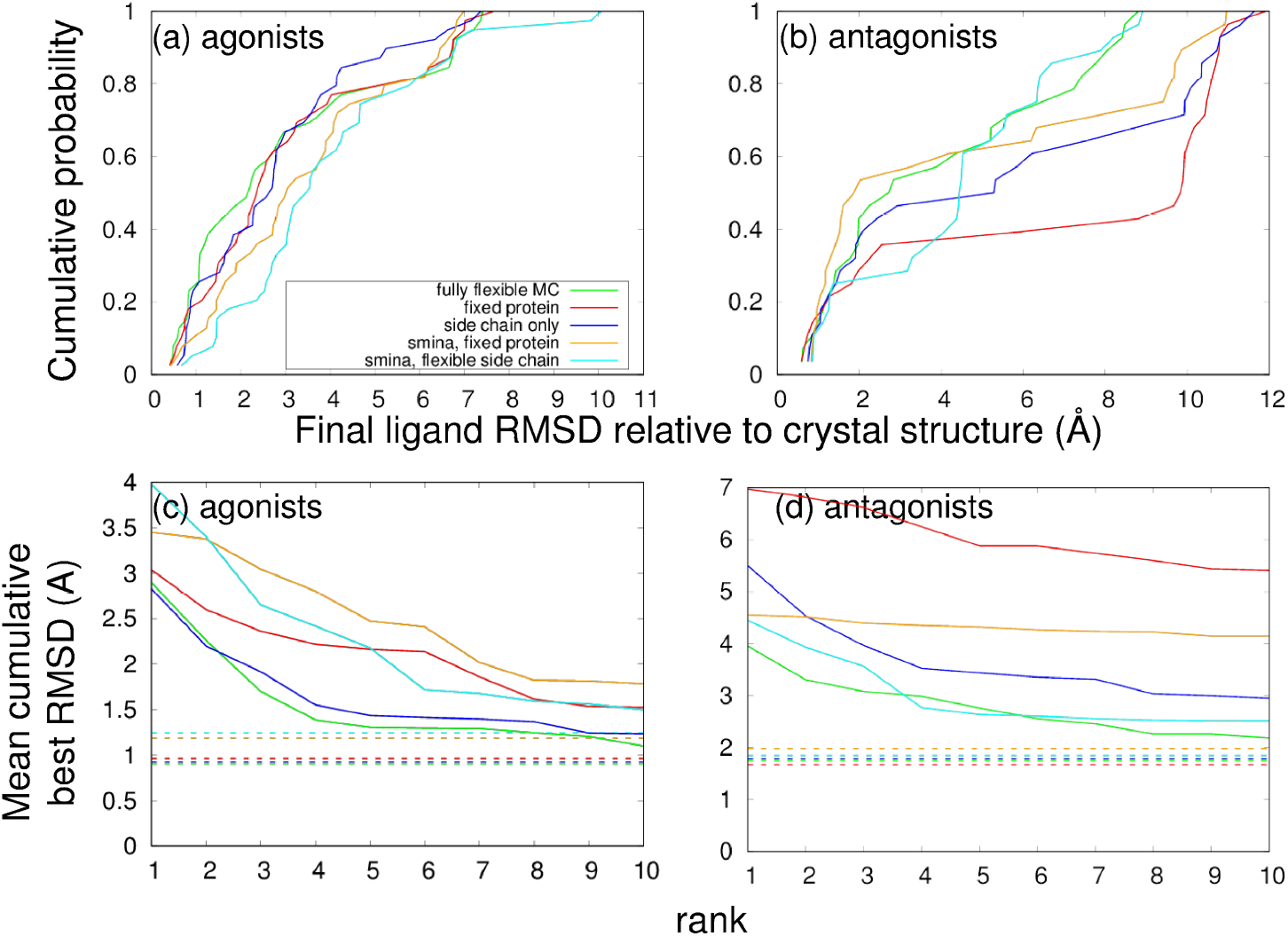
Performance comparison of MRMC with Autodock smina. Plots are similar to those shown in fig. 6. (a) and (b) show the cumulative probability distribution of RMSD values for the docked poses with the lowest final interaction energies for each drug, for (a) agonists or (b) antagonists. Larger values at low RMSD indicate better performance. In (c) and (d), for each drug the docking runs with the *N* lowest interaction energies are chosen and the best RMSD from among these is averaged across (c) agonists or (d) antagonists, so that lower RMSD indicates better performance. This average RMSD is plotted against *N*; dashed horizontal lines indicate the average best RMSD overall, without regard for interaction energy.

The relative computational costs of the two approaches are also of great interest. When run with a fixed protein, Autodock smina performed docking much faster than MRMC. The use of flexible side chains caused smina to slow down considerably; it took 575-650 CPU hours per drug, which is about 1.5-2 times more than MRMC, even though MRMC also includes backbone flexibility in the binding site and CG-based flexibility in the entire protein.

### Assessment of Clustering in Pose Scoring

The fact that some of the docking simulations show substantial changes in the orientation of the ligand relative to the protein suggests that the ensemble of final conformations generated by the MRMC protocol contains information on multiple binding poses which could make a useful contribution to the docking and scoring process. To study this possibility, the ensemble of structures resulting from the docking runs on each drug were divided into clusters based on the protocol described in Sec. Three options for choosing the best conformation from the ensemble were then tested:

1. Based on the observation that, for most drugs, the final interaction energy between protein and ligand correlated with ligand RMSD (Fig. 5), the simplest approach is simply to choose the conformation with the lowest interaction energy, without regard to clusters.
2. Since the largest cluster represents a binding pose that has the highest entropy relative to the other clusters, the conformation within this cluster with the lowest interaction energy could be chosen.
3. Finally, the cluster with the lowest average energy could be chosen, and then the individual conformation with the lowest interaction energy could be chosen from this cluster.

Fig. 9 shows the cumulative distribution of the RMSD of the best conformation for all three of these methods. The differences between them are small, but it appears that method 3 above performs slightly better than the others in that the conformations identified by this method are closer to the crystal structures.

**Fig 9.**
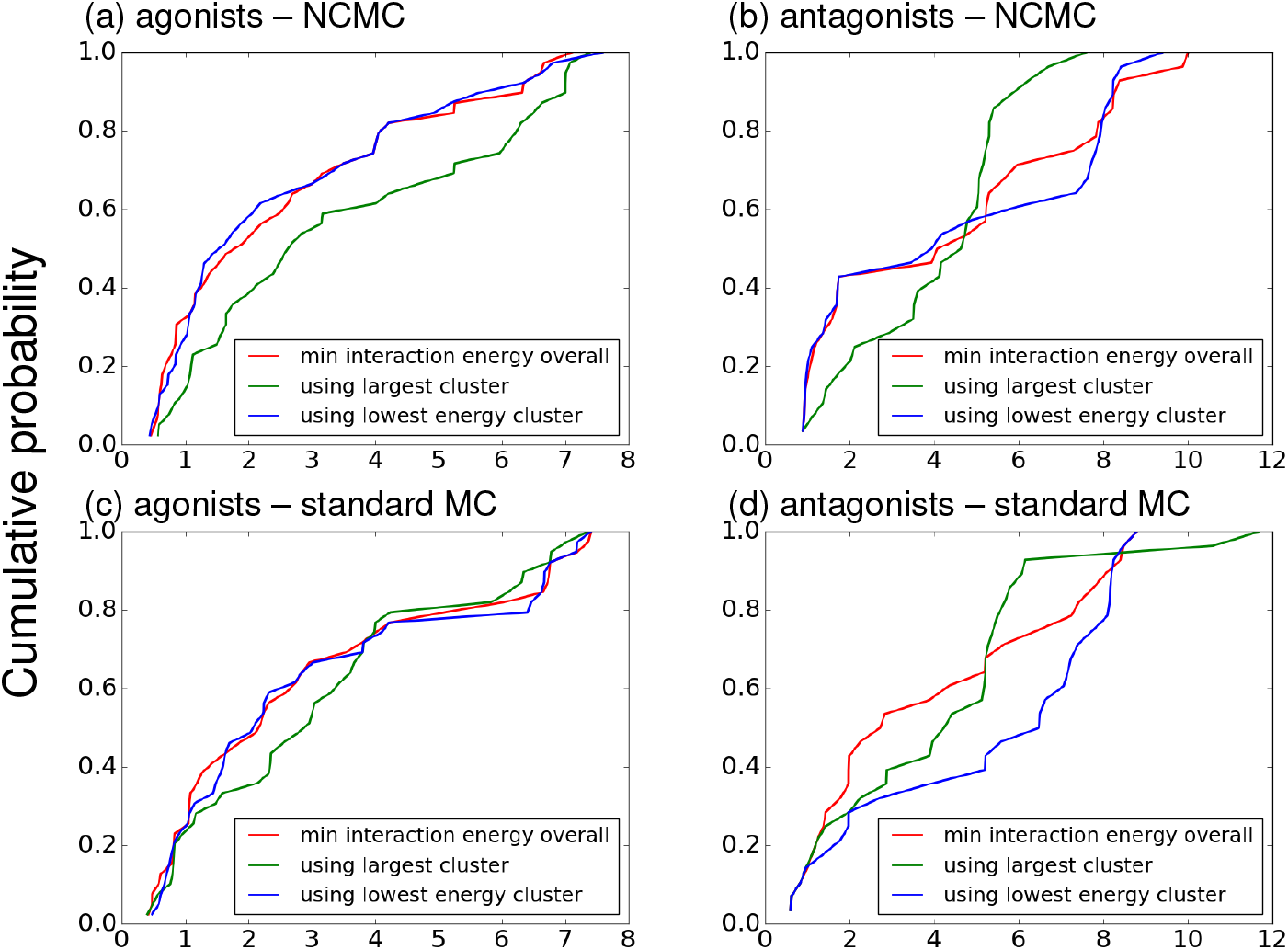
Comparison of methods for ranking ligands. Each graph shows the cumulative distribution of the RMSD of the best conformation (chosen by the indicated method) for (a)-(b) simulations using NCMC or (c)-(d) simulations using standard MC, with a fully flexible protein. (a) and (c) compare ligand-ranking methods for agonists; (b) and (d) do so for antagonists.

Note that we did not employ a more quantitative entropy estimation process because the sampling did not appear to be sufficient - i.e., there were very few jumps between poses in a given MC run (Fig. S2) implying true Boltzmann sampling was not achieved. Also, using the total energy in place of the protein-ligand interaction energy and combining it with similar clustering approaches gave similar results.

### Multiple Poses for WAY-169916

The crystal structure of the ER partial agonist WAY-169916 (PDB code 3OS9, ligand ID KN1) [19] bound to the inactive conformation of ER *α* shows two separate poses for the ligand, with occupancies of approximately 70% and 30%. This provided an opportunity to test whether our protocol can find multiple bound poses for a ligand. The RMSD to each of these poses was calculated separately for our docking simulations of this drug, and a scatterplot is shown in Fig. 10. The ensemble of docked poses for this drug contains two distinct clusters that correspond to the poses in the crystal structure (the closest poses found are approximately 2 Å away from each crystal pose), along with other poses that are similar to neither. This demonstrates that our protocol is able to find both of these poses.

**Fig 10.**
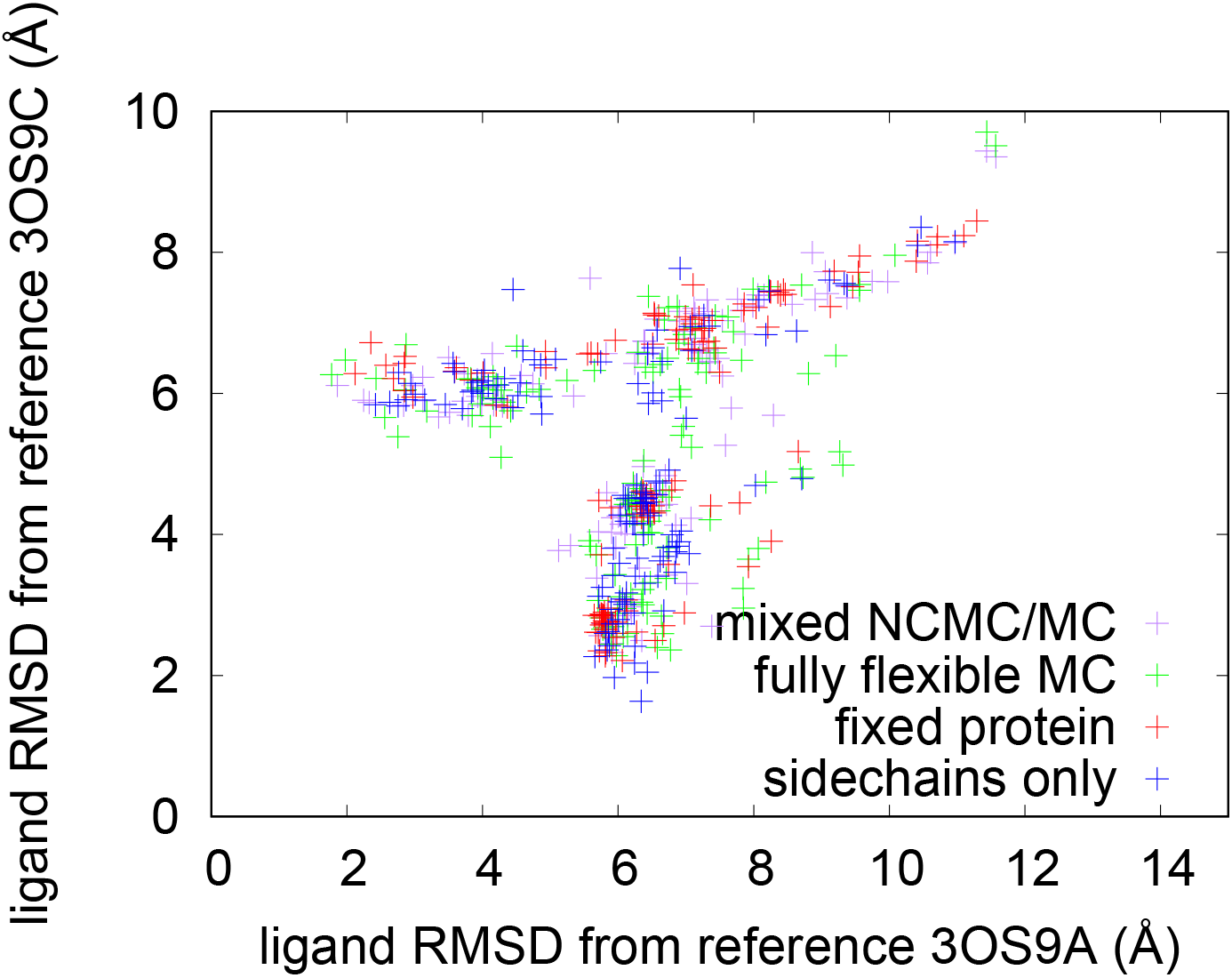
Docking uncovers two crystal poses of the ligand WAY-169916. We show a scatter plot of ligand RMSD relative to the two reference poses found in its reference crystal structure.

### His 524 Ionization States

His 524 plays an important role in the binding of ligands to ER *α*. For example, the O3 atom of estradiol forms a hydrogen bond with a neutral His 524, whereas when 4-hydroxytamoxifen is bound, His 524 instead is ionized and forms a salt bridge with Glu 419. Because of these changes in the ionization state of His 524, and its importance in ligand binding, half of the docking runs were performed with His 524 in an ionized state and half were performed with His 524 in the neutral state. The results for different ionization states of His 524 were generally similar, with approximately equal numbers of successful docking runs (those with a final RMSD less than 2 Å) coming from runs in which His524 was ionized or neutral. Likewise, simulations with both ionized and neutral His 524 produced docked conformations of WAY-169916 that were close to each of the experimental structures.

### Computation time

Table 3 shows a comparison of computation speeds for fully atomistic, mixed-resolution, and coarse-grained representations of ER *α*. As expected, the Gō model is about 60 times faster than a fully atomistic representation. The mixed resolution model is about four times faster than a fully atomistic representation in vacuum, and about twice as fast when the SEDDD solvation model is used. Thus a mixed-resolution model offers a modest savings in compute time, but there is additional sampling benefit from the landscape-smoothing implicitly provided by coarse-graining. As noted, an MRMC platform also offers significant flexibility in implementing docking protocols. For reference, 10 ns of all-atom explicit solvent MD requires about 5 hours with AMBER and one GPU for this system.

**Table 3.** Comparison of computation speeds (MC trial moves per second) for different resolution representations of ER *α* in complex with estradiol.

|  | In vacuum |  | With SEDDD |  |
| --- | --- | --- | --- | --- |
|  | With NCMC | MC only | With NCMC | MC only |
| All atom | 31.69 | 32.52 | 17.70 | 18.39 |
| Mixed resolution | 85.23 | 88.23 | 37.33 | 37.67 |
| Coarse grained only | n/a* | 1279.26 | n/a* | 1279.26 |

\*Coarse grained only results are for uncomplexed ER $\alpha$ ; it is not possible to use NCMC without a ligand.

## Discussion

In this paper, three separate tactics for improving protein-ligand docking were tested. These included a mixed-resolution potential in which most of the protein is treated using a coarse grained model while the region around the ligand was treated atomistically; a nonequilibrium Monte Carlo method, which is intended to improve sampling by systematically varying the coupling between protein and ligand; and the use of clustering to identify free energy basins corresponding to multiple binding poses, and scoring the poses based on this information. The docking results were evaluated by comparing the final structures to known crystal structures; the sampling was also evaluated by studying the relationship between acceptance rate and move size for NCMC moves. We found that allowing for full protein flexibility using the mixed-resolution potential significantly improved the docking results, and the use of NCMC produces a further modest improvement. However, clustering did not appear to offer any significant advantages over simply ranking the poses by protein-ligand interaction energy. Overall, we obtained a correct pose within 2 Å for about half the ligands, so there is significant room for improvement. Below, we discuss ways that several aspects of the approach could be improved. We note that systematic investigation of these different aspects of the docking problem is facilitated by having a highly flexible/adjustable Monte Carlo platform.

### Improving the MRMC Potential

The mixed-resolution model used here is motivated by the concept that the most important approximations in protein-ligand binding will be those between the ligand and the closest amino acid residues within the protein. Therefore, it makes the most sense to model the closest amino acids at the fully atomistic level, while saving computation time by modeling the remainder of the protein using a more approximate coarse-grained model. That said, it is also reasonable to examine every part of the mixed-resolution potential to see if they can be made more physically accurate, while continuing to save computer time over fully atomistic approaches. These include the coarse-grained force field, the atomistic force field, the coupling between them, and the choice of atomistic region.

In this work, a relatively simple Gō model was used to represent the coarse-grained region of ER *α*. This is effective in saving computer time. On the other hand, its reliance on native interactions gives it a strong bias toward the native state, allowing only limited conformational flexibility outside the atomistic region. In the case of ER *α*, the only significant conformational change is the motion of helix 12, so any necessary conformational flexibility could be included by ensuring that helix 12 and the loop connecting it to the rest of the protein were part of the atomistic region. However, with other target proteins this may not be adequate. Additional flexibility could be incorporated by making use of a double well Gō potential [45] or by replacing the Gō potential by another potential that is less dependent on native interactions.

The popular MARTINI force field [77, 78] has been used in a mixed resolution configuration [40] but frequently leads to distorted structures for soluble proteins unless reinforced by an elastic network model. [79] Other potential force fields that could in principle be used include OPEP [80] and UNRES. [81] We have also developed a tunable coarse-grained force field based on constructing interaction energy tables and applying variable amounts of smoothing to them [82]; one of the goals for this force field was to use it in a mixed resolution setting, but substantial additional effort would be required to implement that combination.

Compared to the coarse-grained potential, the atomistic potential used here would seem to offer less room for improvement. The AMBER 99SB force field is a well-tested, commonly used force field, although it could conceivably be replaced with another, newer force field. A more significant area for improvment concerns the treatment of solvation in the simulations. The SEDDD method is based on a distance dependent dielectric with a dielectric constant that includes some solvent exposure. The linear dependence of the dielectric constant on interatomic distance is not physically correct, however, since the dielectric constant should approach that of water as the distance between two atoms increases. There is also no explicit term representing the hydrophobic effect. A generalized Born model [83] would be more physically realistic but also more computationally expensive. Explicit solvent (using water molecules restrained around the atomistic region) would be even better, but would require even more computational cost, and steric clashes between the ligand and water molecules would make it more difficult to sample ligand configurations via Monte Carlo. Another relatively inexpensive solvation model is the Sheffield solvation model, [84] which might provide improved results.

The choice of atomistic region also plays an important role in establishing the tradeoff between physical accuracy and computation time. We selected the atomistic region used here to be as small as possible while including all those residues in direct contact with the ligand in any of the reference structures. We also included helix 12 and its loop to allow for the possibility of transitions between the active and inactive conformations, although we did not observe any such transitions due to the short duration of the simulations. A larger atomistic region would trade computation speed for greater physical accuracy.

### Improved Sampling

While crystal structures of protein-ligand complexes frequently show only one bound pose for the ligand, there is experimental evidence that some ligands can bind to proteins in multiple configurations. The two poses for WAY-169916 are a case in point. Likewise, differences have also been found in ligand binding to aldose reductase depending on the crystallization conditions [18]. In principle, with a sufficient number and length of runs, a docking algorithm based on Monte Carlo or molecular dynamics simulation should be able to find all relevant bound poses for the ligand with the proportion that would be expected based on their relative free energy. Although our MRMC runs indeed found multiple poses for ligands (including both experimental poses for WAY-169916), jumps between poses in a single run were extremely rare.

With our implementation of NCMC, the improvement over standard MC is marginal, in contrast to the success reported recently by the Mobley group for the T4 lysozyme/toluene system using MD. [52] This is likely because the ER *α* ligands studied here are larger than toluene, and shaped in such a way that the barriers separating different ligand orientations are higher than those separating distinct orientations of toluene bound to T4 lysozyme. In addition, our ligands are also more flexible, and we used a larger range of moves and attempted to sample a greater number of degrees of freedom. In an investigation of NCMC applied to side chain rotations of amino acids in explicit solvent, Kurut and coworkers found that NCMC did not enhance sampling for methionine as much as valine, because enhancing the sampling of one dihedral degree of freedom did not improve the sampling of other dihedrals. [85] We may be observing the same effect here, where couplings between the ligand position and orientation and the side chain degrees of freedom of nearby amino acids reduce the efficiency of NCMC. Alternatively, as suggested by Chodera [personal communication] our NCMC protocol may require better targeted ligand MC moves such as “smart darting,” [86] although our initial tests of this idea did not show a substantial improvement.

There are also straightforward means to improve sampling. Two simple ways are to run longer docking simulations or perform more docking runs for each drug. The docking runs used here are fairly short (40000-80000 trial moves) which inherently limits the amount of sampling possible in a single docking run. Of course, both increasing the length and number of docking runs would increase the amount of CPU time needed to dock a drug. In addition, using a longer or otherwise redesigned schedule for *λ*_VDW_ might improve the NCMC acceptance rate, which was relatively low (approximately 2%) in the work reported here, and thereby improve the sampling. Another way to improve the NCMC acceptance rate might be to use a soft core potential for the van der Waals interactions. This might reduce the potential energy changes associated with steric clashes between the ligand and receptor as *λ*_VDW_ is increased from 0 to 1, which contribute to large values of the nonequilibrium work and consequently to poor acceptance rates.

### Use of Clustering Information

If a sufficient degree of sampling can be obtained, it should in principle be possible to identify basins in the free energy surface corresponding to different possible ligand or protein conformations. Each basin will correspond to a cluster of similar conformations obtained in the ensemble. In principle, once all of the basins are identified, if true Boltzmann sampling has been achieved the ensemble of conformations should also give information on the relative free energies of different binding conformations, which can then be used to calculate binding free energies. [87–89] Motivated by this reasoning, we sought to apply clustering algorithms to the ensemble of configurations we obtained from our docking simulations and use this information to aid in identifying the most representative structures. We found, however, that clustering information did not improve the ranking of poses in practice.

The main reason why the clustering was not useful may simply have been that the protocol used here did not allow for adequate sampling, as described above. Another flaw may have been the choice of clustering algorithm or metric used. The complete linkage clustering algorithm used here is relatively crude, being primarily intended for the construction of phylogenetic trees using distances between protein or DNA sequences. [75] Despite this, it was selected because many other algorithms, such as the K-means algorithm, rely on averaging coordinates from distinct configurations, an operation of unclear physical meaning. In addition, complete linkage clustering guarantees that any two configurations classified in the same cluster will have an RMSD less than the selected cutoff. The Cheatham group has tested a number of clustering algorithms on MD trajectories; [90] while they recommend average-linkage hierarchical clustering for circumstances in which the number of clusters is not known in advance (as here) they also point out that the performance of a clustering algorithm is influenced by the choice of atoms used for pairwise comparison and that hierarchical clustering is sensitive to outliers.

## Conclusion

We used a highly adjustable mixed-resolution Monte Carlo (MRMC) platform to examine several aspects of docking protocols in a systematic way. Most importantly, we examined the effects of rigidifying both side chains and the protein backbone. The detrimental results are not completely surprising, but the systematic comparison underscores the importance of backbone flexibility, which is absent from almost all grid-based docking studies. We further examined the sampling improvement afforded by non-equilibrium candidate Monte Carlo, finding only modest improvement in our implementation. Our test case was the flexible ligand binding domain of the estrogen receptor alpha, which is an important cancer target and also a model for other nuclear hormone receptors.

As computing power increases, a ‘middle way’ of docking between grid-based approaches and all-atom free energy calculations may prove useful in drug-design pipelines. This study is a step toward developing such a highly adaptable platform, and already shows improved performance compared to docking software. We recognize that further improvements to sampling and entropy-based pose evaluation will be necessary to make a middle-way tool more valuable for the drug-design enterprise.

## Supporting information

**S1 Agonists used in this work**

**S2 Antagonists used in this work.**

**S1 Analysis of NCMC and MC acceptance probabilities in docking simulations of estradiol to the active conformation of ER *α* as a function of move size.**

**S2 Plots of ligand RMSD relative to crystal structure vs. number of trial moves in docking simulations.**

## Acknowledgments

We thank David Mobley and his group for the suggestion to use the nonequilibrium Monte Carlo method and for valuable discussions. We also acknowledge Andy Stern, John Chodera, David Koes, Jocelyn Sunseri, Ernesto Suarez, Barmak Mostofian and Apoorva Shrivastava for helpful discussions. We also thank the Department of Computational and Systems Biology and the Center for Research Computing at the University of Pittsburgh for computer time. This work was supported by NIH grant no. P41-GM103712, NSF grant nos. MCB-1119091 and CNS-1229064, and a Commonwealth Universal Research Enhancement Program grant from the Commonwealth of Pennsylvania Department of Health (SAP 4100062224).

