## Supplemental Information for "Middle-way flexible docking: Pose prediction using mixed-resolution Monte Carlo in estrogen receptor *α*"

December 23, 2018

| Ligand ID | PDB code for reference structures |
|---|---|
| 0CZ | 3UUA |
| 17M | 2B1Z |
| 1GJ | 4IVW |
| 1GM | 4IU7 |
| 1GQ | 4IUI |
| 1GR | 4IV2 |
| 1GS | 4IV4 |
| 1GT | 4IVY |
| 1GU | 4IW6 |
| 1GV | 4IWC |
| 2OH | 3UU7 |
| 458 | 2B1V |
| 459 | 2FAI |
| 4OH | 3L03 |
| 689 | 1ZKY |
| DES | 3ERD |
| DRQ | 2G5O |
| EED | 2QGT |
| EI1 | 2QAB |
| ESE | 4PPS |
| ESL | 3Q95 |
| EST | 1QKU |
| ETC | 1L2I |
| EZT | 2P15 |
| FSV | 4PPP |
| GEN | 1X7R,2QA8 |
| HZ3 | 2QR9 |
| J2Z | 3HLV |
| J3Z | 3HM1 |
| KN2 | 2QA6 |
| KN3 | 4IW8,3OSA |
| ODE | 2QH6 |
| PIQ | 2QXM |
| STL | 4PP6 |
| T3O | 2G44 |
| WST | 2POG |
| ZTW | 1GWQ |

Table S1: Agonists used in this work.

| Ligand ID | PDB code for reference structures |
|---|---|
| 369 | 3DT3 |
| AEJ | 1XQC |
| AIH | 1XP1 |
| AIJ | 1XP9 |
| AIT | 1XPC |
| AIU | 1XP6 |
| C3D | 2OUZ |
| CM3 | 1YIN |
| CM4 | 1YIM |
| DC8 | 2Q70 |
| E4D | 1SJ0 |
| GW5 | 1R5K |
| I0G | 2I0J |
| IOG | 2IOG |
| IOK | 2IOK |
| JJ3 | 2QE4 |
| KN0 | 3OS8 |
| KN1 | 3OS9* |
| L4G | 2AYR |
| LLB | 2R6W |
| LLC | 2R6Y |
| OHT | 3ERT,2BJ4,2JF9 |
| PTI | 1UOM |
| RAL | 2JFA,2QXS |

Table S2: Antagonists used in this work. *There were two reference structures in this PDB file, which were used separately (see text).

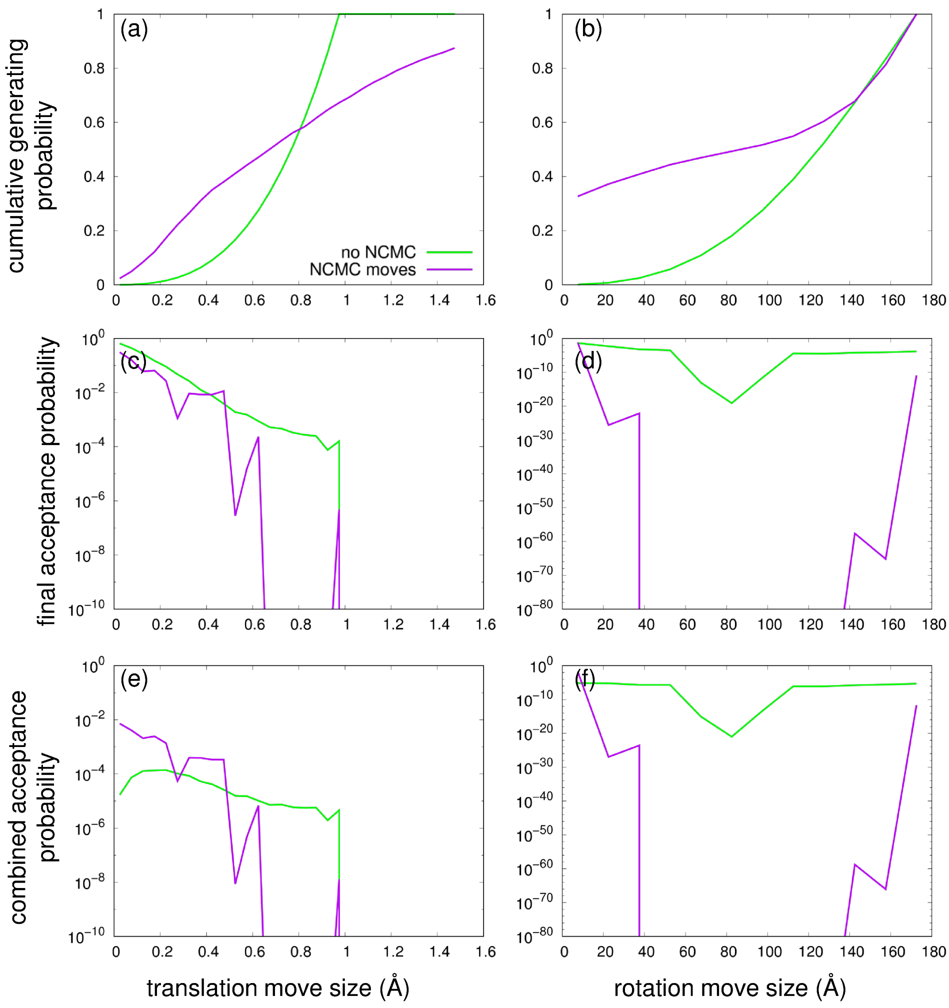


Figure S1: **Analysis of NCMC and MC acceptance probabilities in docking simulations of estradiol to the active conformation of ER $\alpha$ as a function of move size.**
(a), (c), and (e) are for translations; (b), (d), and (f) are for rotations. (a)-(b) Cumulative probability density function of generated move sizes. (c)-(d) Average final acceptance probability of NCMC and MC moves as a function of move size. (e)-(f) Combined acceptance probability (the overall proportion of all NCMC or MC moves that were accepted and of the given size) which is the product of the generating probability.

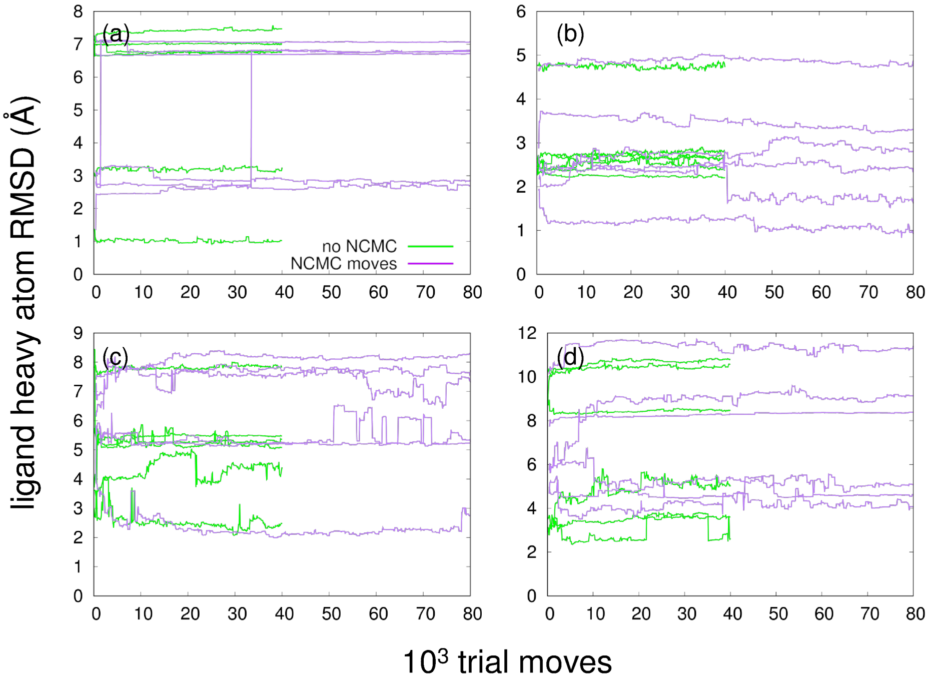


Figure S2: **Plots of ligand RMSD relative to crystal structure vs. number of trial moves in docking simulations.**
(a) genistein; (b) diethylstilbestrol; (c) raloxifene; (d) the drug AIU. Six out of 120 docking runs are shown for each drug.
